# Promoter and transcription factor dynamics tune protein mean and noise strength in a quorum sensing-based feedback synthetic circuit

**DOI:** 10.1101/106229

**Authors:** Yadira Boada, Alejandro Vignoni, Jesús Picó

## Abstract

Gene expression is a fundamental cellular process. Its stochastic fluctuations due to intrinsic and extrinsic sources, known generically as ‘gene expression noise’, trigger both beneficial and harmful consequences for the cell behavior.

Controlling gene expression noise is of interest in many applications in biotechnology, biomedicine and others. Yet, control of the mean expression level is an equally desirable goal. Here, we analyze a gene synthetic network designed to reduce gene expression noise while achieving a desired mean expression level. The circuit combines a negative feedback loop over the gene of interest, and a cell-to-cell communication mechanism based on quorum sensing. We analyze the ability of the circuit to reduce noise as a function of parameters that can be tuned in the wet-lab, and the role quorum sensing plays. Intrinsic noise is generated by the inherent stochasticity of biochemical reactions. On the other hand, extrinsic noise is due to variability in the cell environment and the amounts of cellular components that affect gene expression. We develop a realistic model of the gene synthetic circuit over the population of cells using mass action kinetics and the stochastic Chemical Langevin Equation to include intrinsic noise, with parameters drawn from a distribution to account for extrinsic noise. Stochastic simulations allow us to quantify the mean expression level and noise strength of all species under different scenarios, showing good agreement with system-wide available experimental data of protein abundance and noise in *E. coli*. Our *in silico* experiments reveal significant noise attenuation in gene expression through the interplay between quorum sensing and the negative feedback, allowing control of the mean expression and variance of the protein of interest. These *in silico* conclusions are validated by preliminary experimental results. This gene network could have important implications as a robust protein production system in industrial biotechnology.

**Author Summary:** Controlling gene expression level is of interest in many applications in biotechnology, biomedicine and others. Yet, the stochastic nature of biochemical reactions plays an important role in biological systems, and cannot be disregarded. Gene expression noise resulting from this stochasticity has been studied over the past years both *in vivo*, and *in silico* using mathematical models. Nowadays, synthetic biology approaches allow to design novel biological circuits, drawing on principles elucidated from biology and engineering, for the purpose of decoupled control of mean gene expression and its variance. We propose a gene synthetic circuit with these characteristics, using negative feedback and quorum sensing based cell-to-cell communication to induce population consensus. Our *in silico* analysis using stochastic simulations with a realistic model reveal significant noise attenuation in gene expression through the interplay between quorum sensing and the negative feedback, allowing control of the mean expression and variance of the protein of interest. Preliminary *in vivo* results fully agree with the computational ones.

## Introduction

Noise due to stochastic phenomena is pervasive in the cellular mechanisms underlying gene expression [1, 2]. Its consequences can trigger both detrimental and advantageous effects, and may determine the fate of individual cells and that of a whole population of cells [3–6]. The fluctuations in gene expression of single cells propagate to generate fluctuations in downstream genes affecting stress response, metabolism, development, cell cycle, etc. [5–7], and eventually are the cause of phenotypic noise, that is, variation within an isogenic population.

Experimental measurements of individual genes show that protein production occurs in bursts [4, 8–10] that can be traced back to two main sources: intrinsic, and extrinsic noise. Intrinsic noise is due to stochastic fluctuations in the transcription and translation steps of a particular gene [11]. Thus, intrinsic noise in a gene is correlated with the characteristics of the gene promoter, ribosome binding site (RBS), and the stability of the mRNA and the expressed protein [12, 13]. Transcription dominates the intrinsic noise as the burst size, i.e. the average number of proteins made per mRNA transcript, increases [14]. On the other hand, extrinsic noise corresponds to gene independent fluctuations in protein expression due to external factors like gene copy numbers, transcription factor and ribosome abundance, and/or environmental stimuli [11]. When the gene is encoded on a low-copy plasmid, variability in the gene copy number is a major source of extrinsic noise [15]. This may dominate in eukaryotes [16], while in prokaryotes seems only contribute increasing the noise floor [17].

To minimize the deleterious effects of noise, cells use specific biochemical networks. At the most basic level, cells have evolved different transcription and translation efficiency so as to reduce translation burst rates in key genes [2, 9, 18]. More elaborated strategies, such as negative feedback regulation, may reduce noise by shifting the noise spectrum to a higher frequency region [1, 19]. Ultrasensitive switches and feedforward loops are able to attenuate noise in input signals [20]. These strategies operate at the single-cell level. Yet, cells live in communities, forming a population. At this level, extracellular signaling propagates intracellular stochastic fluctuations across the population [21]. Thus, bacteria have adapted their communication mechanisms in order to improve the signal-to-noise ratio [22]. One of such communication mechanisms is quorum sensing (QS).

Quorum sensing is a cell-to-cell communication mechanism initially discovered in *V. fisheri* and *P. putida* [23, 24]. Bacteria release chemical signaling molecules, called autoinducers, whose external concentration increases as a function of the cell population density. Cells detect a threshold concentration of these autoinducers and alter gene expression accordingly [25–27]. This strategy makes the population as a whole to achieve a desired gene expression level despite the individual noise of each member of the population. It is known that synchronization and consensus protect from noise [28]. Cells consensus induced by diffusion of the signaling autoinducer reduces extrinsic noise by reducing the transmission of fluctuating signals (including noise) in the low-frequency domain [20], and enhances intrinsic stochastic fluctuations [21]. Moreover, quorum sensing allows entrainment of a noisy population when faced to environmental changing signals [29]. Therefore QS seems an effective tool to control the phenotypic variability in a population of cells [30].

Phenotypic variability has important practical relevance in many applications in the areas of biomedicine, biotechnology and other branches of biological science [31]. In particular, the presence of heterogeneous subpopulations may have significant impact on the yield and productivity of industrial cultures [32–34]. Thus, improving homogeneity of protein expression in industrial cultures is a goal of economic relevance for microbial cell factory processes.

Improving homogeneity of protein expression has traditionally been attempted either by optimizing environmental conditions in the culture or by careful selection of the strain. Yet, there is an ever-growing appreciation that biological complexity requires new bioprocess design principles. Synthetic biology, sometimes defined as the engineering of biology, has the potential to engineer genetic circuits to perform new functions for useful purposes in a systematic, predictable, robust, and efficient way [35, 36]. In the last years, several synthetic circuits have been proposed with the ultimate goal of dealing with gene expression noise [20, 22, 37–40].

Synthetic biology makes extensive use of mathematical models and computational simulation to aid the genetic circuit design. They allow the generation of new testable hypotheses and novel ways of intervention, and offer mechanistic explanations of experimental results. The dynamics of the reactions involved in gene expression have been traditionally described using continuous deterministic mathematical models [41, 42]. These equations can be easily obtained from the reactions using the law of mass action kinetics [43]. However, this continuous deterministic approach fails to capture the consequences of stochasticity in gene expression. Hence, there is an extensive literature on rigorous modeling of gene expression noise and determination of its sources [44].

The most accurate way to represent the stochasticity originated from intrinsic sources is by means of the Chemical Master Equation (CME). The CME determines the probability that each species will have a specified number of molecules at each time instant [44, 45]. Though the CME can be solved analytically for small systems [46, 47], it is in general not a tractable problem, or even possible, for systems of medium to large size. Not even numerically. Several alternatives have been derived with a broad range of computational cost and precision. Some numerical schemes like those in [48, 49] provide a solution of the CME in a truncated state-space. However, long-term predictions are not always possible, specially for bimolecular reactions. In [50] the authors present a modified CME based on the partition of a system into reactions with low and high propensity, thus providing approximations at different levels of accuracy and computational cost. The Gillespie SSA algorithm [51] is a widely used Monte Carlo-based approach that provides exact samples from the probability distribution that results from solving the CME. However, having an interconnected population of cells, as in our proposed gene synthetic circuit, jeopardizes the possibility of employing SSA for several reasons. First there are different volumes involved, extracellular and intracellular. The diffusion through the membrane of the autoinducer molecule used for cell-to-cell communication depends on its concentration in both of them, making the account for the probability of reaction more complicated. Second, when using SSA, several realizations or trajectories of the system are needed in order to obtain an accurate estimation of the statistical moments of the species in the circuit, making the use of SSA in a population of interconnected cells a computationally very demanding task. At the opposite extreme the linear noise approximation (LNA) allows to get the decoupled dynamics of the mean and variance of gene expression [19, 52, 53] using a first order approximation (i.e. a linearization) of the difference between the actual noisy trajectory, and the mean deterministic one. An intermediate practical alternative to model gene expression intrinsic noise is the use of the Chemical Langevin Equation (CLE).

The CLE is a stochastic differential equation (SDE) driven by zero-mean Gaussian noise that describes the system when the molecules of reactants into a cell population are sufficiently large [54, 55]. It approximates the CME by a system of stochastic differential equations of order equal to the number of species –*cf*. order equal to the possible number of molecules of all the species in the CME. Extrinsic noise can be modeled by randomizing values of the model parameters [16, 56], an approach that can easily be integrated within the CLE framework. The CLE has also been used to study intrinsic noise in synthetic systems involving QS mechanisms in [20] though the authors considered an averaged cell, thus not taking into account single cell contributions to the noise in the population.

In this work we analyze a gene synthetic network designed to reduce gene expression noise while achieving a desired mean expression level. The circuit combines a negative feedback loop over the gene of interest, and a cell-to-cell communication mechanism based on quorum sensing. In the Materials and Methods Section, we develop a realistic model of the gene synthetic network over the population of cells using mass action kinetics. Then, stochastic simulations using the Chemical Langevin Equation allow us to quantify the noise strength of all species under different scenarios. In particular, in the Results Section we analyze the ability of the circuit to reduce noise as a function of parameters that can be tuned in the wet-lab, and the role quorum sensing plays. We show preliminary experimental data validating our *in silico* results, and finally we draw some conclusions in the Discussion Section.

## Materials and Methods

### Description of the synthetic gene network

Reducing gene expression noise at the level of an individual cell can be attempted in several ways. Open loop strategies as based on sensitivity analysis providing guides as to how properly tune transcriptional and translational parameters so that the noise levels can be controlled while the mean values can be simultaneously adjusted to desired values [57]. While sensitivity analysis gives very valuable insights, open loop control is not robust against system uncertainty and/or variations. On the other hand, closed loop control can be implemented by using a negative feedback loop over the gene of interest and appropriately tuning it. Though negative feedback has been proved to decrease gene expression noise [37], a single-cell intracellular feedback loops do not take into account that in practice one is interested in controlling gene expression mean value and noise across a population of cells. Feedback across a population of cells can be implemented by means of quorum sensing-based strategies, and has been shown to reduce noise effects [20, 22, 30]. Indeed, cell-to-cell communication by means of quorum sensing induces consensus among cells [58], that is contributes to reduce the difference of internal state among cells in a population. This, in turn, contributes to protect from noise [28]. Thus, the idea of joining both intracellular negative feedback and extracellular feedback via quorum sensing is a natural one, that has been suggested in [38, 39].

Here we analyze the gene synthetic network proposed in [38], designed to reduce gene expression noise while achieving a desired mean expression level. The circuit combines a negative feedback loop over the gene of interest, and a cell-to-cell communication mechanism based on quorum sensing. For that purpose, the circuit employs two functional subsystems already implemented in *E. coli* (see Fig 1A).

**Figure 1.**
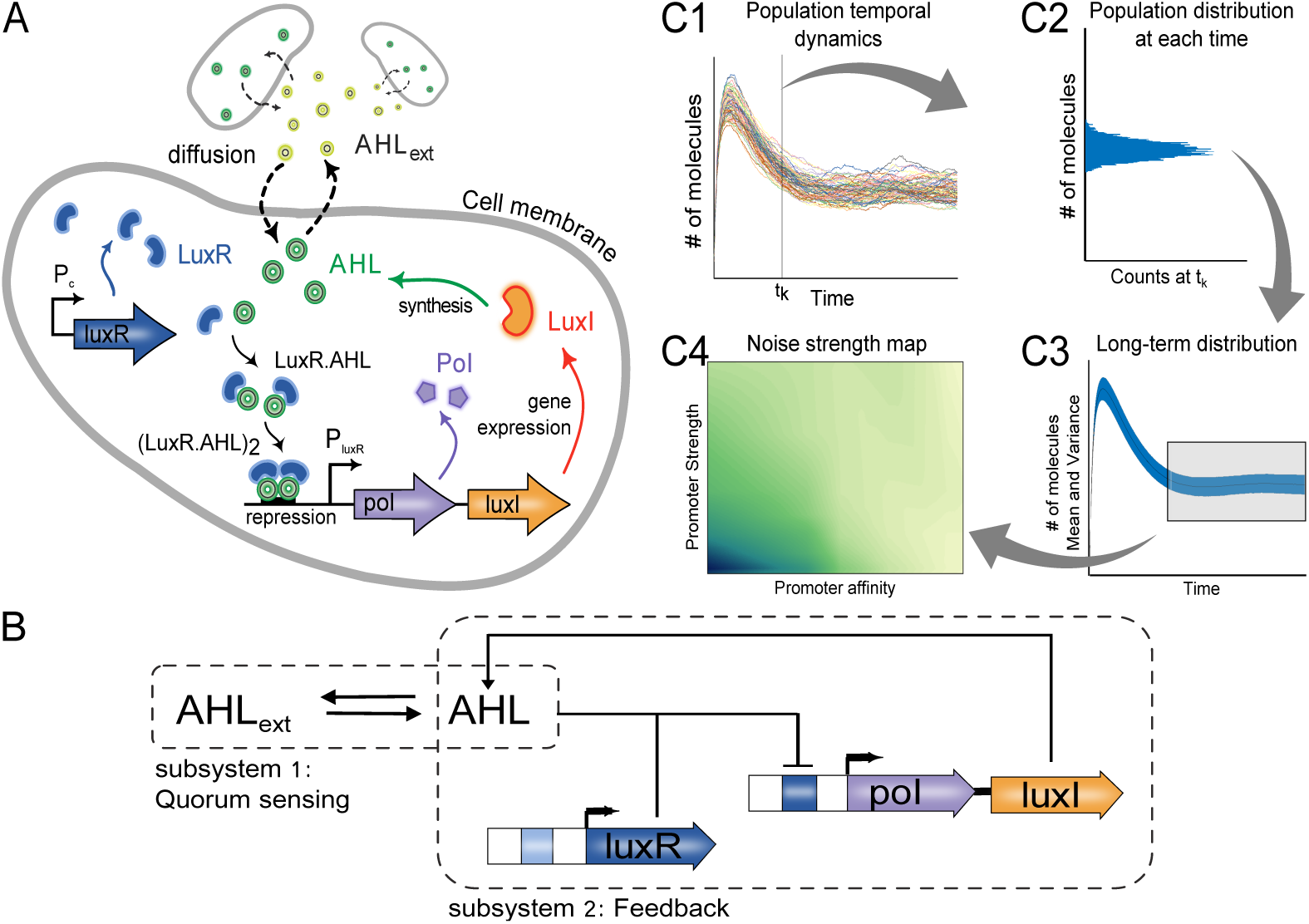
Synthetic gene network. **A.** Intracellular and extracellular system with negative feedback control and quorum sensing mechanism. **B.** Biochemical reactions and diffusion process. **C.** Methodological procedure to obtain the statistical moments from stochastic simulations of the circuit. C1) Temporal evolution of one species in the population of cells. C2) Distribution of the number of molecules across the population at each time instant. C3) Acquisition of the long-term distribution for each species. C4) Noise strength map for varying model parameters.

The first subsystem implements a cell-to-cell communication mechanism via quorum sensing, based on exchange of the small signaling autoinducer molecule N-acyl-L-homoserine lactone (AHL) [25, 26]. This autoinducer molecule passively diffuses across the cellular membrane to and from the external environment. Intracellular AHL is synthesized by the protein LuxI expressed by an homolog of the gene luxI of *V. fisheri* [59].

The second subsystem uses the synthetic repressible promoter P_luxR_ designed in [60] to control transcription of gene luxI. This promoter is repressed by the transcription factor (LuxR.AHL)_2_. Protein LuxR is expressed by gene *luxR* under the constitutive promoter P_c_. Proteins LuxR and AHL bind forming the heterodimer (LuxR.AHL), which subsequently dimerizes forming the heterotetramer (LuxR.AHL)_2_. This way a negative feedback control of the LuxI expression is effectively implemented [38].

The circuit acts as a closed loop controller of the mean and variance of a protein of interest. This protein can be either fused to protein LuxI, or coexpressed with it. In the first case, a linker is inserted between the fused proteins allowing intracellular self-cleavage using a TEV protease [61–63]. Alternatively, if the protein of interest is coexpressed with LuxI, the controller will only act at the transcriptional level. In cases where transcriptional noise dominates translational one, e.g. when the average number of proteins made per mRNA transcript is larger than two [14], coexpressing LuxI with the protein of interest is a simple yet effective approach.

**Note:** Henceforth, and for the sake of simplicity, we will call *monomer* the heterodimer (LuxR · AHL), and *dimer* the heterotetramer transcription factor (LuxR · AHL)_2_.

### Mathematical model

In order to analyze how our genetic circuit affects intrinsic and extrinsic noise we need an appropriate model, and a computationally efficient method. Both aspects are intertwined. We use the Chemical Langevin Equation approach. Though computationally much more efficient than the CME or even the Gillespie algorithm, the CLE is still computationally demanding when the goal is to simulate a whole population of cells. Since the CLE approximates the CME by a system of stochastic differential equations of order equal to the number of species, a reduced model with as few species per cell as possible is desirable. Thus, in a first step we use the mass-action kinetics formalism [42, 43] to get a deterministic model of the full reactions network corresponding to the genetic circuit. We then get a reduced order model by applying the *Quasi Steady-State Approximation* (QSSA) on the fast chemical reactions and taking into account invariant moieties [53, 64, 65]. We aim at obtaining a reduced model more amenable for computational analysis, but avoiding excessive reduction that would lead to lack of biological relevance. In particular, the species we obtain in the reduced are not lumped ones. Reduced models accounting for total mRNA and total transcription factor have been proposed to match modeled species with measurable ones [66]. In our case we explicitly model bound and unbound forms of the transcription factor, but the model accounts for the total LuxI protein. For our circuit this is a good proxy of the amount of protein of interest if both are co-expressed, and transcriptional noise dominates. In the best case, when the protein of interest is in self-cleavable tandem fusion with LuxI, both will express in 1:1 stoichiometric ratio [62]. Moreover, the resulting lumped parameters in the reduced model are easy to associate to tuning knobs available in the wet-lab implementation in the relevant cases [67], and their values are amenable to be obtained experimentally.

In a second step we use the deterministic reduced model to infer a stochastic CLE-based model whose mean corresponds to that of the deterministic one.

#### Reduced deterministic model

We start from considering the relevant biochemical reactions in the circuit. We consider the *gene expression* sets of reactions on the one hand, and the *induction* ones on the other. In the *gene expression* block, the main processes considered for each of the proteins are transcription, translation, mRNA degradation and protein degradation. In the *induction* block, the reactions considered are reversible binding between the protein LuxR and the inducer AHL to form the monomer (LuxR.AHL), monomer degradation, reversible dimer (LuxR.AHL)_2_ formation and its degradation, diffusion of the inducer across the cell membrane, extracellular and intracellular inducer degradation, and reversible binding of the dimer to the repressible promoter P_luxR_. We assumed the promoter P_luxR_ is leaky.

From the resulting set of reactions, and based on mass-action kinetics we derived a complete deterministic model using ordinary differential equations (ODE) for number of molecules of each species. We considered the net effective transcription rates of genes *luxI* and *luxR*, and took into account the basal transcription rate (leakage) of the repressible promoter P_luxR_. Besides, the dilution effect due to the cells growth rate was added to every degradation rate. This complete model was then reduced using QSSA, conservation of moieties, and assuming that translation, diffusion across the cell membrane, and dimerization are the dominant dynamics. With this assumptions we got the dynamic ODE model:

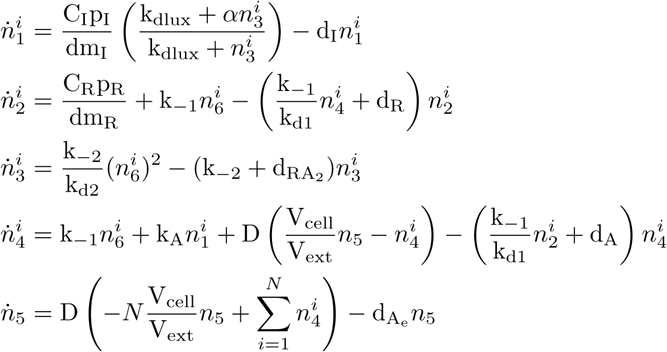

with:

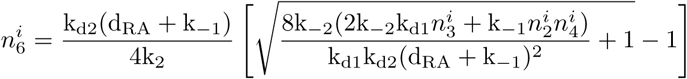

where 
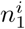
 is the number of molecules of protein LuxI in the i-th cell, 
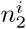
 that of LuxR, 
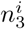
 is the dimer repressor (LuxR.AHL)_2_, 
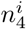
 is the intracellular amount of inducer AHL, *n*_5_ is the external one, and 
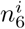
 is the amount of monomer (LuxR.AHL) molecules. Notice this is given as an algebraic equation obtained assuming that monomerization is faster than dimerization, and can be assumed to be at quasi-steady state, as confirmed by the comparison between the complete and the reduced model dynamics. Table 1 describes the parameters in the model and its nominal values.

**Table 1.**
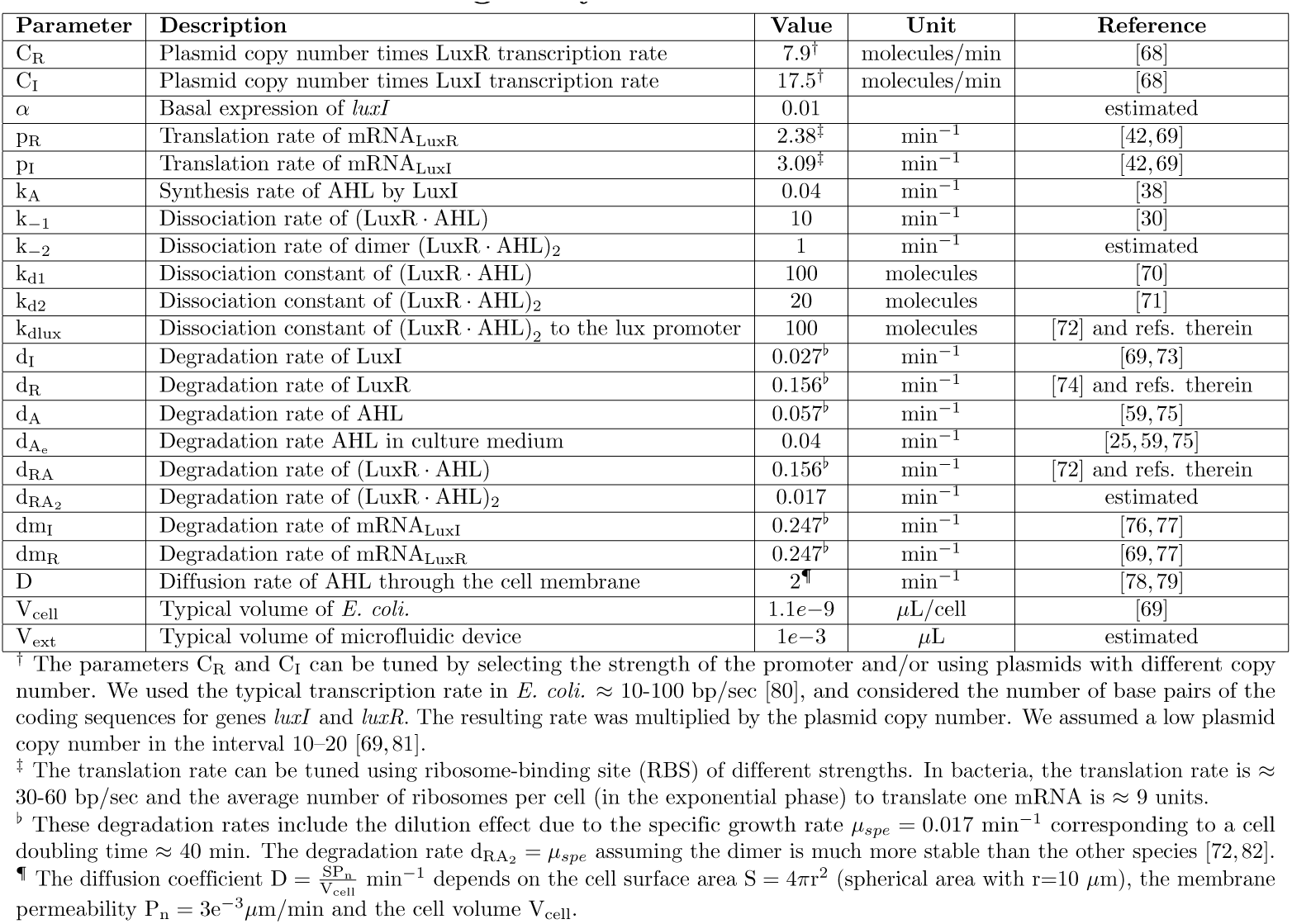
Parameters of the gene synthetic circuit model.

#### From the deterministic ODE model to the stochastic CLE one

To model gene expression intrinsic noise we derive a stochastic CLE-based model whose mean corresponds to that of the deterministic model (1)-(2). This can be done by considering an equivalent set of pseudo-reactions for the deterministic model. From these, one can set model (3), corresponding to the Euler-Maruyama discretization of the CLE for a system with a population of *N* cells:

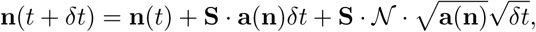

where ***n***(*t*) = [***n***(*t*)^*i*^,…*n*(*t*)^N^, *n*_5_]^*T*^ are the number of molecules of each species in the population, being ***n***(*t*)^*i*^ the vector of species LuxI, LuxR, (LuxR.AHL)_2_, and intracellular AHL for the *i*^th^ cell, and *n*_5_ the extracellular one AHL_ext_. The stoichiometry matrix **S**, whose elements are the stoichiometric matrices of each cell **S**_cell_ and the external stoichiometry **S**_ext_, has structure:

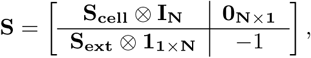

where ⊗ is the Kronecker product, **I**_**N**_ the identity matrix of dimension *N* × *N*, **0**_**N** × 1_ and **1**_1 × **N**_ are vectors of zeroes and ones respectively, and matrices **S**_**cell**_ and **S**_**ext**_ are expressed by equations (5).

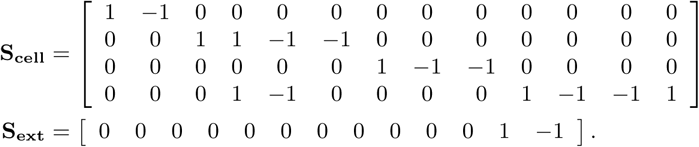

The coefficients in the stoichiometric matrices (5) were obtained from the equivalent set of pseudo-reactions for the deterministic model, and the term **a**(**n**) in model (3) is the associated vector of reaction propensities for the whole population of cells, with:

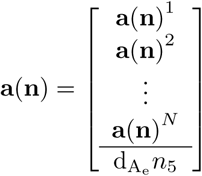

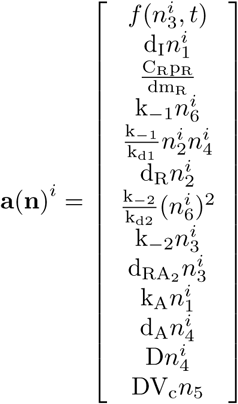

where 
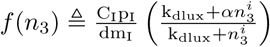
 is the Hill-like function associated to LuxI repression.

Finally, 
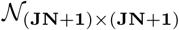
 where *J* = 13 is the number of reactions for the *i*_th_ cell, i.e. the dimension of (7), is a diagonal matrix with continuous normal random variables of zero mean and unit variance as elements.

Notice we use lumped propensity functions in (7) derived from the reduced model, like the *f*(*n*_3_) Hill-like function associated to LuxI repression. This approach has already been used in [83]. We validated it for our model by simulating the pseudo-reaction associated to *f*(*n*_3_) using CLE, and comparing the result with that obtained by simulating the set of corresponding original reactions using Gillespie’s direct method SSA.

Extrinsic noise can be modeled by randomizing the values of the model parameters [16, 56], an approach that can easily be integrated within the CLE framework. In particular, we assumed a normal distribution in the model parameters to account for the extrinsic noise.

### Computational analysis

We use the stochastic model (3) of the proposed circuit, hereafter denoted as circuit Qs/Fb, to explore the impact of some key circuit parameters on noise. As control circuit to compare with, we consider a second circuit which removes both QS and the feedback loop, denoted as NoQs/NoFb. For the computational analysis, this accounts to setting the synthesis of AHL to zero (*k*_*A*_ = 0[1/min]) in model (3). This condition is achieved in the lab experimental implementation by taking out the gene coding for LuxI (see). To asses the effect of cell-to-cell communication, we also considered a hypothetical circuit with feedback but without quorum sensing (NoQS/Fb, *D* = 0). Notice the circuit NoQS/Fb cannot actually be implemented for it assumes there is no diffusion of the autoinducer molecule across the cell membrane. Yet, it is useful as a computational thought experiment to account for the contribution of the cell-to-cell communication.

Gene expression noise is evaluated using the squared coefficient of variation, i.e. the noise strength measure (*η* ^2^ = (*σ*/*μ*)^2^). The noise strength measure *η*^2^ properly captures the contributions of both intrinsic and extrinsic noise [84], and allows comparisons for different expression rates.

We followed the general procedure depicted in (Fig 1C). First, for different combinations of the model parameters, we performed simulations of the temporal evolution of the number of molecules of each species in the circuit for each cell in the population involved in our system (Fig 1C1). Extrinsic noise was modeled by randomizing the values of the model parameters using a normal distribution with a variance of 15%. The models were implemented using OpenFPM^1^, a C++ version of the Parallel Particle Mesh (PPM) library allowing efficient computational particle-mesh simulations [85]. In all simulations we used a population of *N* = 240 cells in a culture volume of 10^−3^ *μ*l, corresponding to an optical cell density OD_600_ = 0.3. Cell density variations did not appreciably change the results, confirming the results in [20].

Then, we obtained the first two statistical moments *μ* and *σ*^2^ for each species in the cell population at every time *t*_*k*_ (Fig 1C2). We used the laws of total expectation and total variance. Using these moments, we calculated long-term distributions to infer the noise strength of each species (Fig 1C3). To this end, we checked with our models that one realization of the population of *N* cells is enough to obtain unbiased values of the long-term moments of the population, provided there is enough time to perform the time average.

Finally, we generated noise strength maps for different sets of varying model parameters (Fig 1C4). We explored the effect of variations in parameters associated to expression of LuxI and LuxR, as they are as key parameters in our circuit. For LuxI we considered the dissociation constant k_dlux_ between the transcription factor (LuxR · AHL)_2_ and the repressible P_lux_ promoter, the translation rate p_I_, and the basal expression *α*_I_ of the P_lux_ promoter. We sampled in the ranges k_dlux_ = [10 – 2000], *α* = [0.01 − 0.1], and p_I_ = [0.2 − 10] selected from the literature [86–88] and experimentally achievable in the lab. As for LuxR, we considered two values for the the translation rate p_R_: a strong RBS (p_R_ = 10), and a medium-weak one (p_R_ = 2). In addition, we analyzed the effect of different degradation rates d_R_ in the range [0.02 – 0.2].

Notice from model (1) that although we only considered variations in the translation rates p_I_ and p_R_, these are tantamount to considering variations in the lumped values 
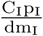
,

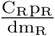
 corresponding to the products of protein burst size, transcription rate and gene copy number. We assumed variations in translation rates just because they are relatively simple to modify in a graded way by tuning the RBS [86], though also transcription rates could be easily tuned [89].

### Plasmids and experimental conditions

To validate the *in silico* computational results, we implemented the Qs/Fb and NQs/NFb circuits *in vivo*. We used components from the iGEM Registry of Standard Biological Parts (http://parts.igem.org). All parts were cloned using the Biobrick’s foundation 3 Antibiotic Assembly method. All coding sequences have the double-terminator BBa_B0015, and were confirmed by sequencing.

The circuit Qs/Fb integrating both the QS-based cell-to-cell communication and the negative feedback subsystems, was split in two subunits integrated in different plasmids. On the one hand, plasmid pCB2tc contains the gene *luxR* (part BBa_C0062) coding for the protein LuxR constitutively expressed under the control of a medium strength promoter (part BBa_J23106) and a strong RBS (part BBa_B0034). This insert was cloned into the pACYC184 plasmid cloning vector (p15A origin, chloramphenicol/tetracycline). On the other hand, plasmid pYB06ta contains *gene luxI* (part BBa_C0161) under control of the p*_LuxR_* repressible promoter (part BBa_R0062) and a strong RBS (part BBa_B0034). The strong RBS BBa_B0034 and the green fluorescent protein (GFP, part BBa_E0040) were inserted using GIBSON assembly (NEB Catalog# E2611S) upstream of *luxI*, right after the p*_LuxR_* promoter. This way, GFP, used as protein of interest (PoI in Fig 1) is co-expressed with *LuxI*. They were inserted into the pBR322 plasmid cloning vector (pMB1 origin, ampicillin/tetracycline). Finally, both plasmids pCB2tc and pYB06ta were co-transformed in competent cells (DH-5*α*, Invitrogen).

As control circuit, we implemented the circuit NQs/NFb which removes both QS and the feedback loop. To this end, the plasmid pCB2tc above was co-transformed with the plasmid pAV02ta (pMB1 origin, ampicillin/tetracycline) containing only GFP downstream the p*_LuxR_* repressible promoter (part BBa_R0062) and the the strong RBS (part BBa_B0034). Both were cloned in the pBR322 plasmid cloning vector.

For the experimental validation of the circuit (see protocol details in Supplementary Text S3), two sets of *E. coli* cells (cloning strain DH-5*α*) carrying the Qs/Fb and NoQs/NoFb circuits respectively were inoculated from −80°C stocks into 3 mL of LB with appropriate antibiotics, followed by an overnight incubation at 37°C and 250 rpm in 14 ml culture tubes. When the cultures reached an optical density (OD) of 4 (600 nm, Eppendorf BioPhotometer D30), the overnight cultures were diluted 500-fold (OD of 0.02) into M9 medium with appropriate antibiotics. These were used to inoculate new cultures, which were incubated for 7 hours (37°C, 250 rpm,14 ml culture tubes) until they reached an OD between 0.2–0.3. At this point, cell growth and protein expression were interrupted by transferring the culture into an ice-water bath for 10 min. Next, 50 *µ*L of each tube were transferred into 1 ml of phosphate-buffered saline with 500 *µ*g/mL of the transcription inhibitor rifampicin (PBS + Rif) in one 5 mL cytometer tube, and incubated during 1 hour in a water bath at 37°C, so that transcription kept blocked and GFP had time to mature and fold properly. Samples were measured at different time points using the BD FACSCalibur flow cytometer (original default configuration parameters), with and without adding AHL_e_ as external disturbance (10nM N-3-Oxohexanoyl-L-homoserine lactone, Santa Cruz Biotecnology Catalog Number SC205396).

## Results

### Quorum sensing and negative feedback attenuate gene expression noise

We first addressed the question whether the proposed QS/Fb circuit effectively reduces noise strength with respect to the circuit NoQS/NoFb. Recall the last one consists of the LuxR expression on the one hand, and the protein of interest (PoI) downstream the p*_LuxR_* repressible promoter, without the luxI gene coding for LuxI protein, on the other (see Fig 2A). Since no autoinducer *AHL* is neither produced nor externally introduced, there is no repression, so expression of PoI is essentially a constitutive one. This corresponds to the Poisson distribution observed in the population histograms in Fig 2B. Contrarily, the QS/Fb histogram departs from the Poisson distribution to become a narrow Gaussian-like one. This fact, and the reduction in the mean expression value, indicate the strong presence of regulation. In both cases we used the nominal parameters (see Table 1).

**Figure 2.**
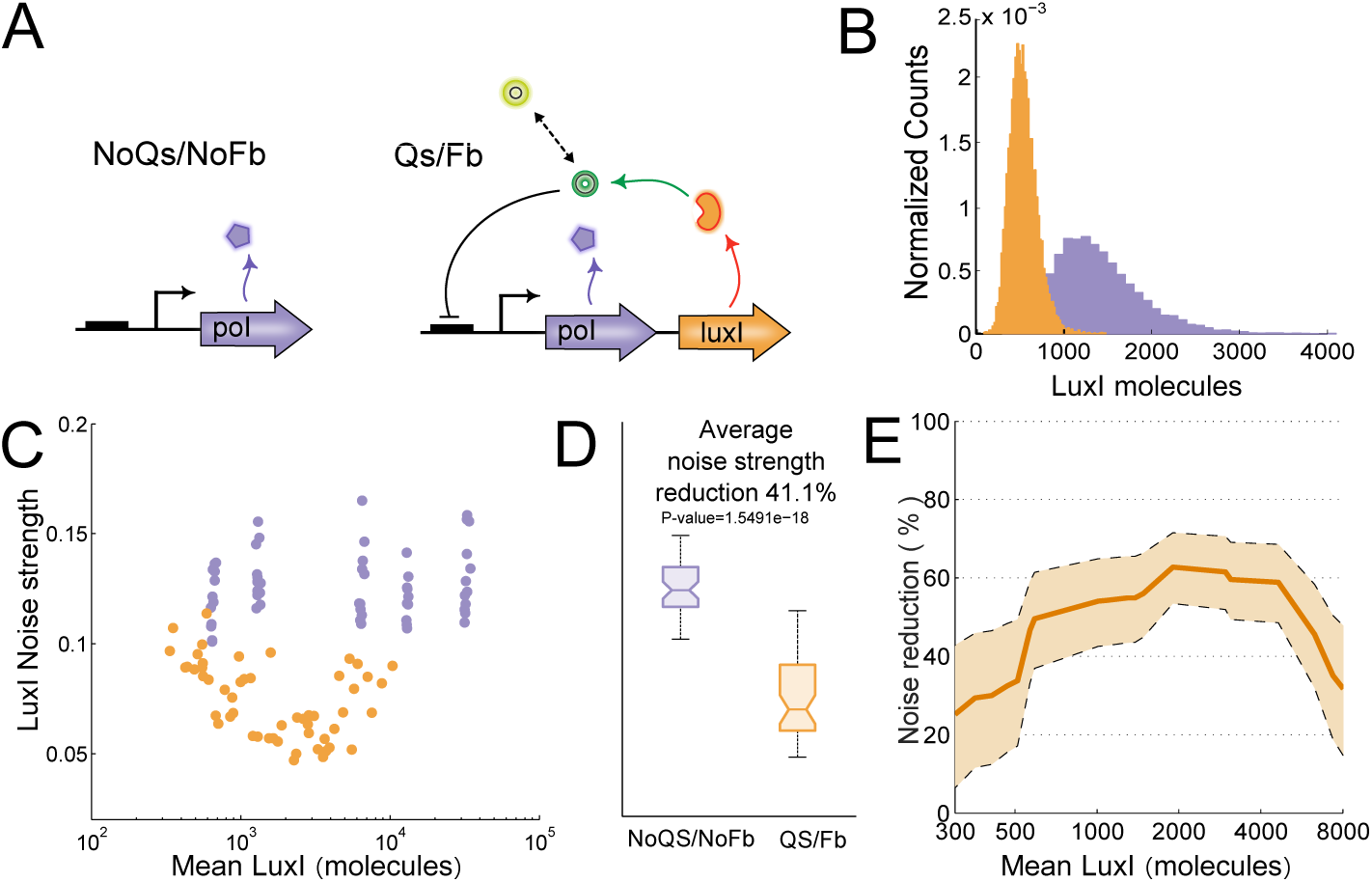
LuxI noise strength under presence/absence of quorum sensing and negative feedback. A. Circuits topologies: NoQS/NoFb (top) and QS/Fb (bottom). B. Representative computational population histograms of LuxI noise strength for QS/Fb (orange) presenting a narrower gaussian-like distribution as compared to the Poisson-like one of NoQS/NoFb (purple). C. Sampled combinations of LuxI expression characteristics for fixed LuxR ones show larger values of LuxI noise strength *versus* mean for NoQS/NoFb (purple dots) than for QS/Fb (orange dots) D. The QS/Fb circuit significantly reduces the average noise strength for the sampled parameters space by 41%, from 〈*η*^2^_NoQS/NoFb_〉 = 0:1263 down to 〈*η*^2^_QS_/_Fb_〉 = 0.0744. E. For varying LuxR parameters the average reduction of noise strength in LuxI ranges from 30 % up to 60 % and shows dependence on the mean expression level.

Reduction in noise strength was not due to a particular choice of the circuit parameters, but a property of the proposed topology. Fig 2C depicts LuxI noise strength *versus* mean expression for 60 different combinations of the p*_LuxR_* repressible promoter characteristics (see methods section) for both Qs/Fb (orange points) and NoQs/NoFb (purple points). The points in the figure correspond to the mean values across the cells population for each parameters combination. The magnitude of the noise strength reduction was larger for medium values of mean protein expression. Noise strength levels were similar for all mean expression values in the case of the NoQs/NoFb circuit. Mean expression values in this case depend only on the translation rate p_I_ for which five discrete values were used, inducing the five mean values seen in the figure. On the contrary, the Qs/Fb circuit showed lower values of noise strength and more graded values of the mean expression level, for it depends on the combination of all three parameters varied.

More importantly, the noise strength was consistently lower for the Qs/Fb circuit. Taking together all the different combinations of promoter parameters for each circuit, the average noise strength was significantly reduced by 41% in the presence of quorum sensing and negative feedback, decreasing from
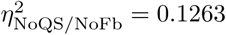
 down to 
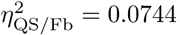
, as shown in Fig 2D.

For given fixed LuxR expression parameters, the noise strength reduction in LuxI showed a clear dependence on its mean expression level. To elucidate if this dependence was only due to the choice of the p*_LuxR_* promoter parameters we evaluated, for each mean expression level, the ratio between the noise strength of the Qs/Fb circuit for all the range of p*_LuxR_* promoter parameters, and that of the NoQs/NoFb circuit. With this, we obtained the results plotted in Fig 2E showing the minimum and maximum values of LuxI noise variance reduction as a function of its mean value. In the range between between 600 and 6000 LuxI molecules it was possible to reduce the noise variance at least in 35% in the worst case scenario, with a maximum reduction of around 70% for means between 2000 and 3000 molecules.

### Feedback pays-off when extrinsic noise dominates

At this point the question arises as to what are the roles of quorum sensing and that of feedback in noise strength reduction, and what are their effect in view of both intrinsic and extrinsic noise.

To answer this question we first contextualized the computational results using available experimental data of noise strength and protein abundance in *E. coli.* We used experimental data taken from [90], and plotted it against our computational results in three scenarios: base control circuit with no quorum sensing nor feedback (NoQS/NoFb, *k*_*A*_ = 0), our circuit with both quorum sensing and feedback (QS/Fb), and the hypothetical circuit with feedback but without quorum sensing (NoQS/Fb, *D* = 0). For each scenario we considered different combinations of parameters under the same conditions as in section, with values of the mean protein number in the range 10^0^ − 10^5^.

Fig 3A shows the experimental data plotted as black dots. The dashed red and blue lines are the intrinsic and extrinsic noise limits respectively, taken from the same reference. Simulations including both intrinsic and extrinsic noise are plotted as purple dots (NoQS/NoFb), green (NoQS/Fb) and orange ones (QS/Fb) using the same data as in Fig 2C. Our computational results fully agreed with the experimental data and derived limits in [90]. The results corresponding to the base control circuit NoQS/NoFb clearly were over the noise limits.

**Figure 3.**
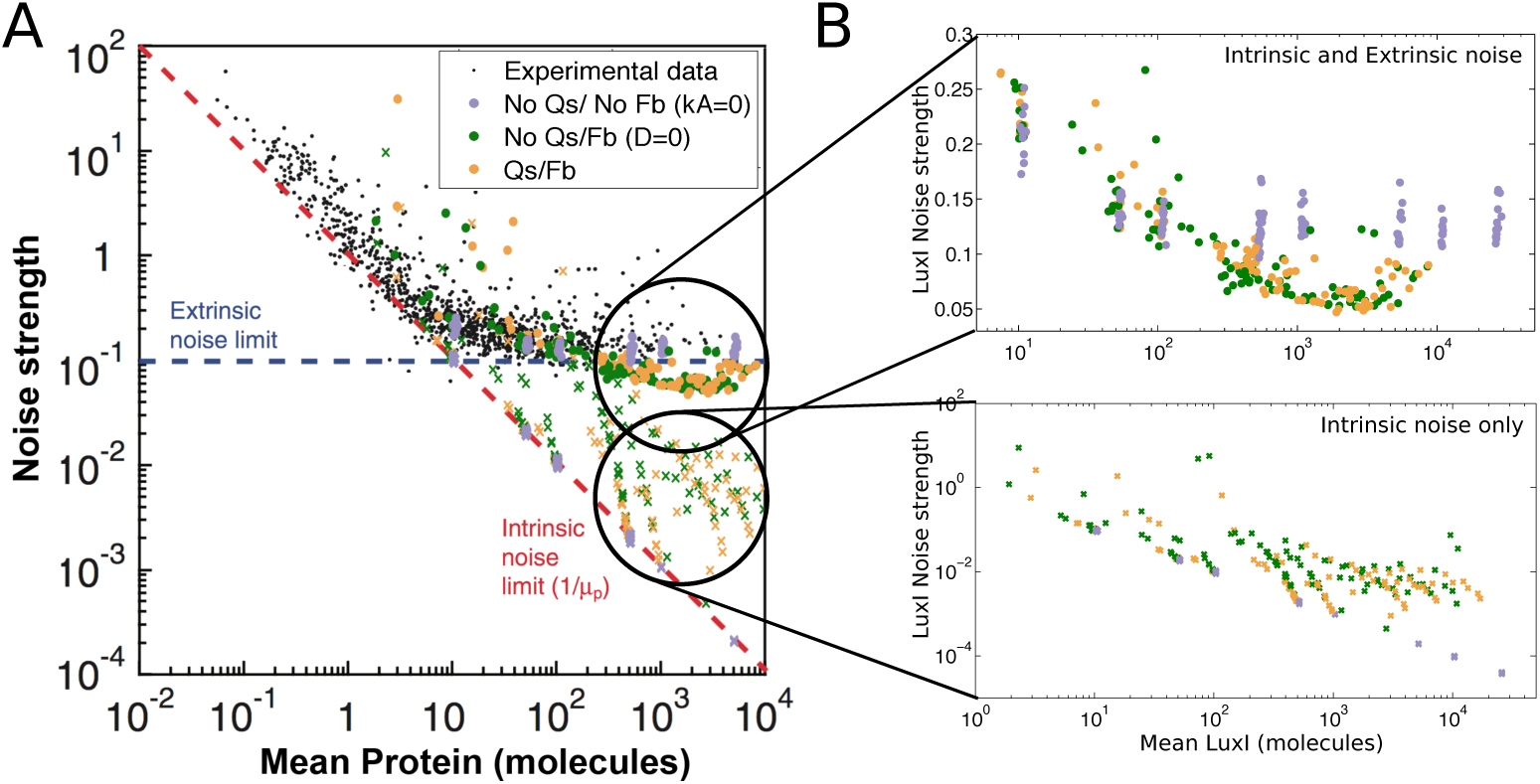
Comparison between experimental data and different scenarios evaluated computationally. A. Experimental data of protein abundance and noise in *E. coli* taken from [90] is plotted as black dots. The dashed red and blue lines are the intrinsic noise limit and the extrinsic noise limits respectively, taken from the same reference. Simulations of the gene circuits in our study, including both intrinsic and extrinsic noise, are plotted using purple dots (NoQS/NoFb), green (NoQS/Fb) and orange ones (QS/Fb). Simulations including only intrinsic noise are plotted as crosses: violet (NoQS/NoFb), green (NoQS/Fb) and orange (QS/Fb). B. Zoom of the scenarios considering both intrinsic and extrinsic noise (top) and only intrinsic noise (bottom).

Unexpectedly, noise strength of both circuits QS/Fb and NoQS/Fb integrating feedback showed very similar behavior. As shown in Fig 3B the QS/Fb and NoQS/Fb points lay in the same region. For medium and high mean protein expression values noise strength in QS/Fb and NoQS/Fb decreased just below the reported extrinsic noise limit, and well below the noise strength for the base NoQS/NoFb circuit. Though high protein expression are of main interest for the intended application of our circuit in an industrial biotechnological context of heterologous protein production, we were interested in the performance of the circuits at low mean protein numbers. Interestingly, the situation in the region was reversed. The open loop circuit NoQS/NoFb showed consistent lower noise strength values than QS/Fb and NoQS/Fb. Therefore, feedback contributed to reducing noise strength for medium-high protein expression where extrinsic noise dominates.

### Quorum sensing helps feedback to cope with intrinsic noise

The last result was inconclusive about the contribution of quorum sensing to reduce noise strength. To settle this issue we concentrated our analysis in the medium-high protein expression region where feedback contributed to reduce noise strength and extrinsic noise dominates.

We first wanted to elucidate whether QS mainly contributed reducing the intrinsic component of noise. If this was the case, its effect could be masked by the dominant extrinsic noise. To that end we carried out simulations for the same combinations of parameters as before, but suppressing extrinsic noise, and considering the three scenarios NoQS/NoFb, QS/Fb, and NoQS/Fb. The results are shown in Fig 3A, plotted as violet (NoQS/NoFb, *k_A_* = 0), green (NoQS/Fb, *D* = 0) and orange crosses (QS/Fb). Fig 3B shows a zoom into the relevant region. Introducing either feedback alone or feedback plus quorum sensing increased noise strength values with respect to the minimal base control circuit representing plain constitutive protein expression. The results for this base NoQS/NoFb circuit were along the experimental intrinsic noise limit derived in [90]. These results were consistent with the findings at low mean protein values where intrinsic noise dominates. The circuit NoQS/Fb with feedback and no cell-to-cell communication showed higher values of noise strength, specially for lower values of mean protein number. Finally, reintroducing quorum sensing (QS/Fb) was able to slightly improve noise strength.

To confirm this result we evaluated the difference between the noise strength in LuxI between the circuits QS/Fb and NoQS/Fb when only intrinsic noise is present as a function of circuit parameters associated to LuxI expression. Fig 4(A) shows the noise strength map difference for different combinations of the dissociation constant k_dlux_ vs. the LuxI translation rate p_I_ when we consider a tight promoter P_lux_, *α* = 0.01 or a leaky one *α* = 0.1 in both noise scenarios. The noise strength reduction when QS was added reached a 200% for low values of p_I_. Increasing the dissociation constant improved the reduction, specially for a leaky promoter.

**Figure 4.**
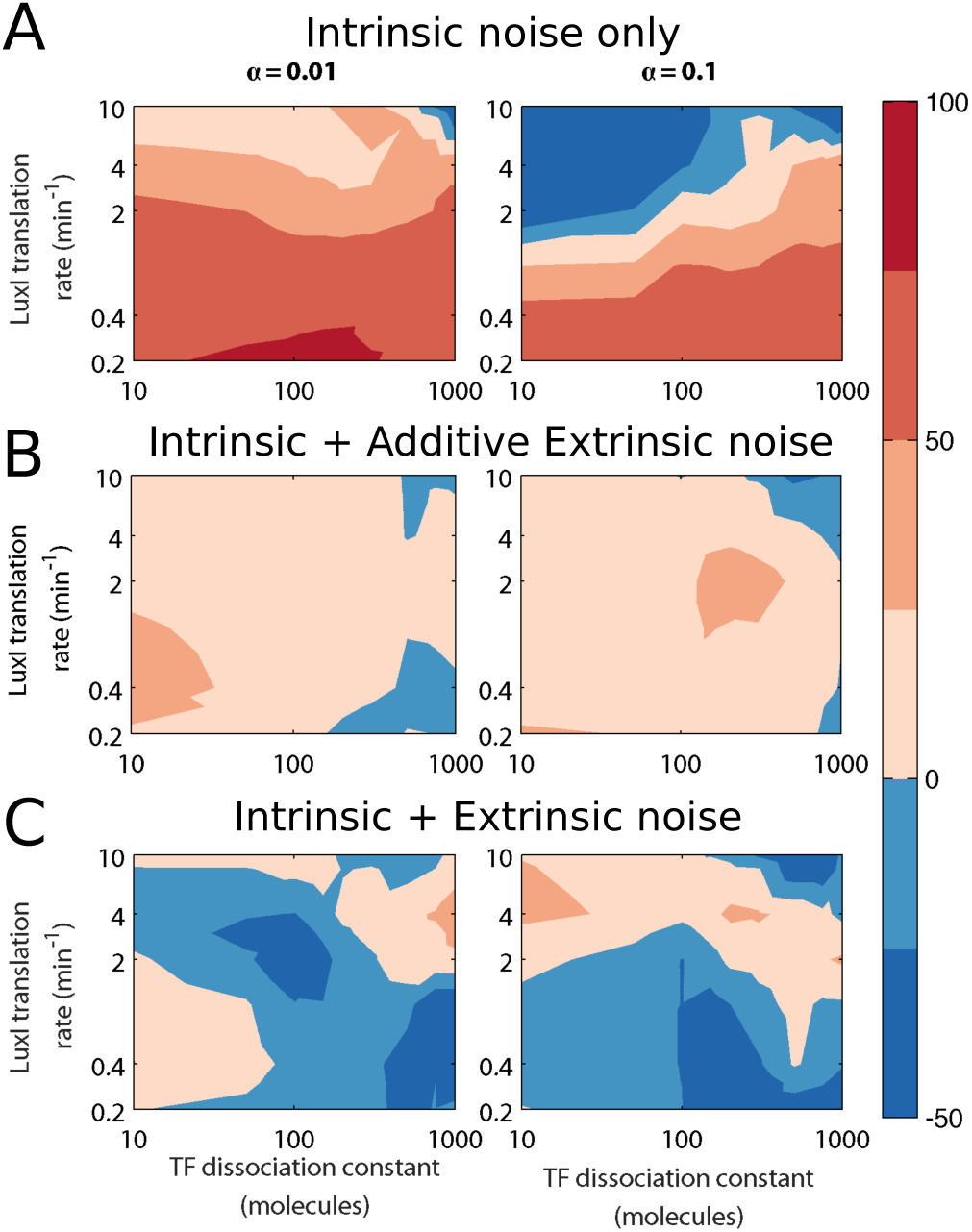
LuxI noise strength reduction as a function of circuit parameters. Color map of the reduction of LuxI noise strength when QS is added to Fb w.r.t. the dissociation constant k_dlux_ and the LuxI translation rate p_I_. All other parameters were set to their nominal values. Left) Tight promoter *α* = 0.01. Right) Leaky promoter *α* = 0.1. A) Only intrinsic noise is present. B) Intrinsic noise is present incorporating also additive extrinsic noise. C) Intrinsic and extrinsic noise present.

The previous result suggested that the results reported in the literature showing a reduction in noise strength when QS was used were a result of modeling extrinsic noise as an additive signal. This hypothesis was confirmed when besides intrinsic noise we introduced an additive extrinsic noise to our system, with variance independent of the system states. Fig 4(B) shows that in this case there also was a generalized noise strength reduction for most parameter combinations.

Finally, in case we restored extrinsic noise as parametric variability the results showed that adding QS may increase or decrease noise strength (Fig 4(C)) strongly depending on the values of the circuits parameters, and suggesting that getting benefit of QS for medium-large mean expression values requires optimizing the circuit parameters tuning.

### Tuning LuxI expression allows minimising noise-strength

Dependence of mean expression and noise strength on the Qs/Fb circuit parameters is a key factor to understand for it to be of potential practical usage. To this end we performed thorough *in silico* experiments to estimate the noise strength and mean expression value of LuxI, as a proxy of the protein of interest, for different sets of the circuit parameters associated to LuxI expression as described in the methods section. We only evaluated two values for the basal expression, corresponding to a tight P_lux_ promoter (*α* = 0.01), and a leaky one (*α* = 0.1). As for LuxR, we also considered two values: a strong RBS (p_R_ = 10), and a medium-weak one (p_R_ = 2). We kept all other parameters to their nominal values described in table 1. Recall that although we considered variations in the translation rates p_I_ and p_R_, these are tantamount to considering equivalent variations in the lumped values of the corresponding products of protein burst size, transcription rate and gene copy number.

Fig 5 shows the noise strength map for different combinations of the dissociation constant k_dlux_ vs. the LuxI translation rate p_I_ when we consider both a tight promoter P_lux_, *α* = 0.01 (Fig 5A) and or a leaky one *α* = 0.1 (Fig 5B). The means of LuxI protein number are shown as contour lines.

**Figure 5.**
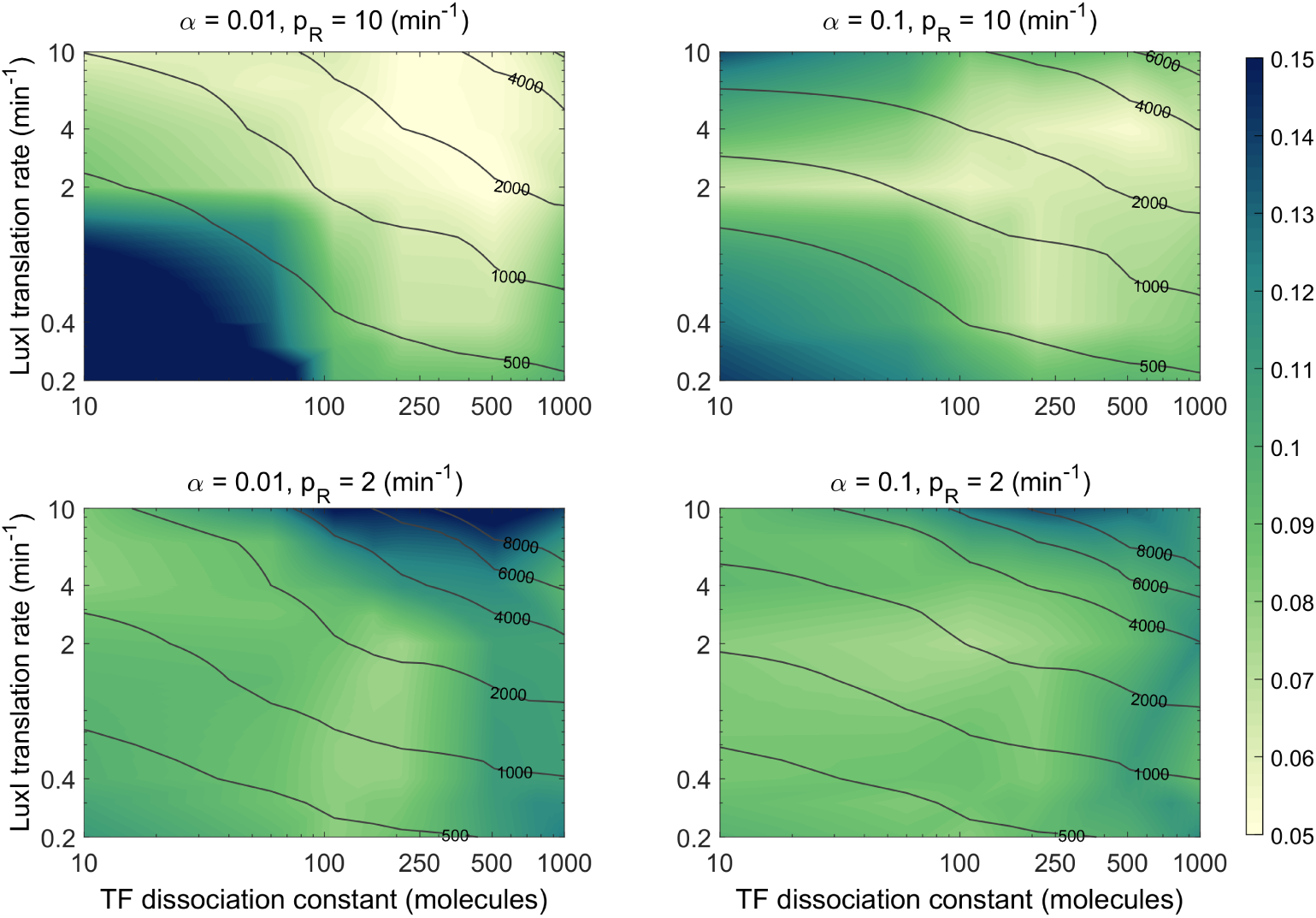
LuxI noise strength and mean as a function of circuit parameters. Color map of LuxI noise strength w.r.t. the dissociation constant k_dlux_ and the LuxI translation rate p_I_. The level curves correspond to the mean number of LuxI molecules. A)Strong LuxR RBS with p_R_ = 10 [1/min]. B) Medium-weak LuxR RBS with p_R_ = 2 [1/min].

The mean expression levels of LuxI followed general monotonous trends in all cases. It increased for simultaneous increasing of the dissociation constant and the LuxI translation rate. On the other hand, increasing leakiness of the LuxI promoter did tend to lower mean expression levels of LuxI for low values of the dissociation constant. Finally, using a weaker RBS controlling the translation of LuxR (Fig 5B) produced a steeper increasing of the mean expression level as the dissociation constant and the LuxI translation rate increase.

Noise strength did not show simple patters as a function of the circuit parameters. Larger variations between high and low noise strength values were observed for stronger LuxR RBS (Fig 5A) independent of the leakiness of the promoter P_lux_. In this case, the lowest values of noise strength were achieved for values of the dissociation constant k_dlux_ in the range [100 – 500], and values of LuxI translation rate p_I_ in the range [2 – 10]. The mean expression levels in this region were between 2·10^3^ and 4·10^3^ proteins, in agreement with the results shown in Fig 2. Decreasing the LuxR RBS strength kept the the values of minimal noise strength essentially in the same region, but with higher values (Fig 5B). The same trend towards higher values of noise strength was observed when the tight promoter P_lux_ was changed for a leaky one. This was more evident when a stronger LuxR RBS was used (Fig 5A).

### Fast LuxR turnover reduces LuxI noise strength

Next we analyzed in more detail the effect of LuxR expression parameters on LuxI mean expression level and noise strength. In particular, we were interested in the effect of the LuxR translation rate *p_R_*, as main tuning knob of the LuxI mean expression level, and the one of the degradation rate *d_R_*, as it has been suggested that fast LuxR turnover can reduce the extrinsic noise. We fixed the LuxI traslation rate to two values *p_I_* = 1 and *p_I_* = 2 [min^−1^] around its nominal value, and again considered both a tight pLux promoter (*α* = 0.01) and a a leaky one (*α* = 0.1). We kept all other parameters to their nominal values described in table 1.

Fig 6 shows the LuxI noise strength maps and mean expression level curves as a function of values of the LuxR translation rate in the range 0.2 to 10 1/min, and LuxR degradation rate in the range 0.02 to 0.2 1/min. The mean expression level did depend little on the LuxR degradation rate, with a slight increase for large ones. LuxR translation rates or, tantamount, LuxR synthesis rates, proved to be a good sensitive tool to tune the desired LuxI mean expression rate, with larger values of the last as the former decreased.

**Figure 6.**
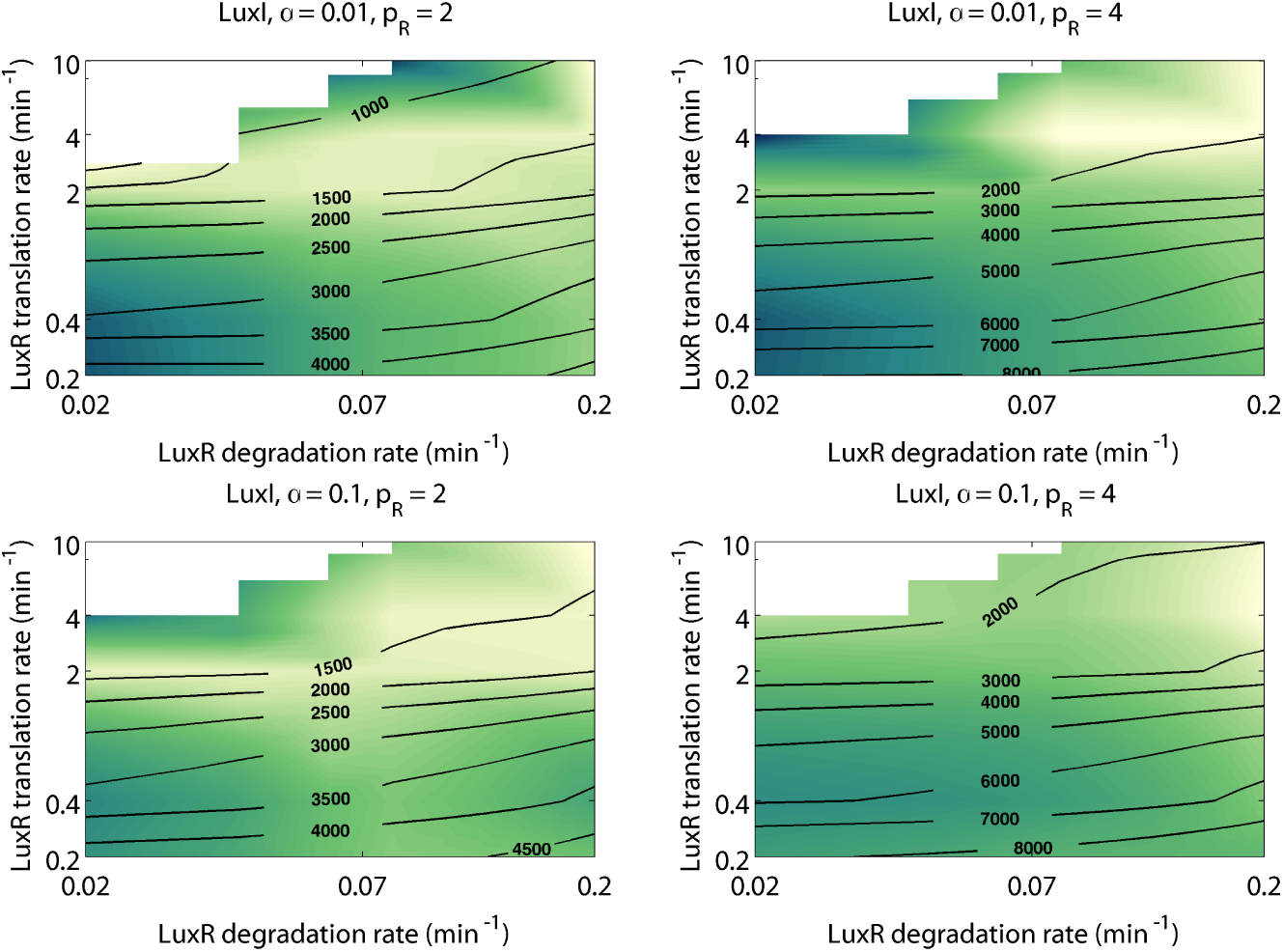
LuxI noise strength *versus* LuxR parameters. LuxI noise strength maps and mean expression level curves for a tight pLux promoter (*α* = 0.01, top) and a a leaky one (*α* = 0.1, bottom) with LuxI translation rates *p_I_* = 2 (left) and *p_I_* = 4 (right) around its nominal value.

Interestingly, LuxI noise strength decreased as LuxR degradation rate increased, with optimal values in the range 0.07 to 0.2 1/min (*cf*. nominal value in table 1). It did not show a clear dependence on LuxR translation rate, but for an interesting region, for values of LuxR around its nominal value, where it tended to decrease for all values of the degradation rate.

### Experimental results confirm computational predictions

Experimental implementation of the proposed QS/Fb circuit would not only allow a preliminary experimental validation of its capability to reduce noise strength, but would also further validate the model parameters used throughout this study. Recall the comparison in Fig 3 referred our results to the general landscape of mean protein abundances and noise strengths in *E coli.* To this end we experimentally implemented both the NoQS/NoFb and QS/Fb circuits as described in section, and compared the experimental results with the ones obtained using the corresponding nominal computational models.

The steady state population histograms of LuxI for the circuits Qs/Fb (orange) and NoQs/NoFb (purple) under the same experimental conditions are depicted in Fig 7. The computational predictions for the nominal models are in the panel A, while the B panel shows flow cytometry experimental results. Both results were qualitatively comparable without any tuning, fitting or change in the model parameters. We only required a common scaling factor to convert from relative units of fluorescence to number of molecules. The experimental results showed LuxI noise strength is reduced by 31.5% 
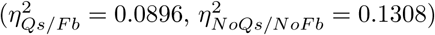
. On the other hand, the computational simulations predicted a 33.6% reduction 
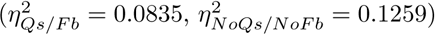
.

**Figure 7.**
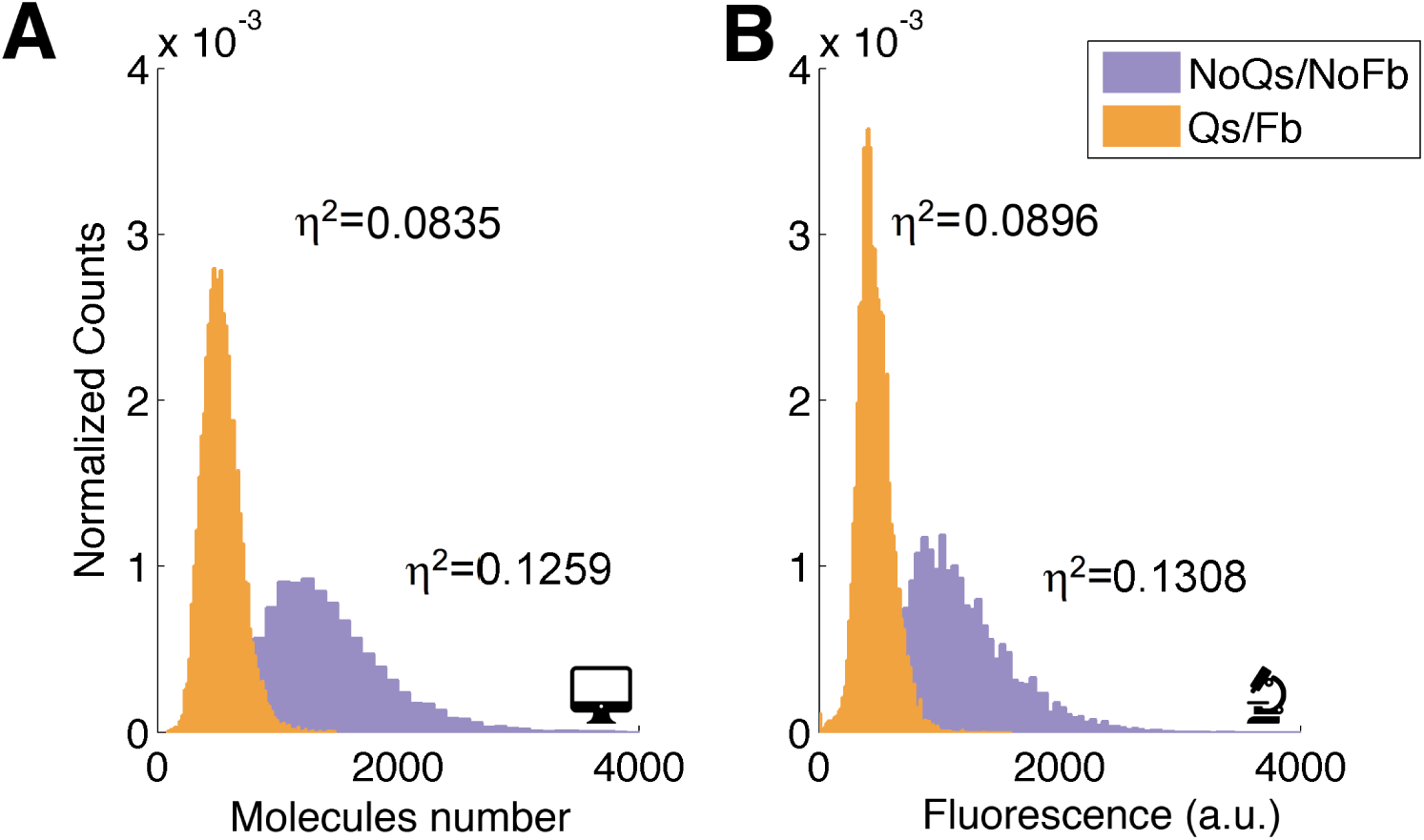
Comparison between experimental and computational results. Population distributions and noise strength for both Qs/Fb and NoQs/NoFb circuits. The plots in A correspond to the computational models simulation, and the ones in B correspond to experimental flow cytometry results.

As we expected, the mean expression level in the NoQs/NoFb circuit, *μ_NoQs/NoFb_* = 1166.46 (a.u.) (simulation *μ_NoQs/NoFb_* = 1395.65 (molecules)), was larger than the one in the Qs/Fb circuit, *μ_Qs/Fb_* = 462.63 (a.u.) (simulation *μ_Qs/Fb_* = 533.67 (molecules)). Indeed, the circuit Qs/Fb incorporates a feedback loop that changes the location of equilibrium point with respect to the open loop circuit. To test whether the reduction in noise strength is an effect associated with the reduction in the mean expression level, we compared both circuits applying an induction of 3nM AHL to the the open loop NoQs/NoFb circuit. With this induction, GFP fluorescence mean values of both circuits was comparable.

## Discussion

Our results show that gene synthetic circuits benefiting from the interplay between feedback and cell-to-cell communication allow control of the mean expression level and noise strength of a protein of interest. A few circuit parameters easy to tune in the wet-lab can be used to achieve noise strength reductions up to a 60% with respect to constitutive expression of the protein of interest.

Mean expression level and noise strength are not independent goals. At low mean values intrinsic noise dominates and sets the minimum noise strength attainable. At high mean values extrinsic noise dominates. Thus, there is a trade-off between expression level and noise strength, as revealed both by system-wide experimental data and theoretical analysis reported in the literature. Our computational results fitted well in this scenario, and suggest that tuning synthetic gene circuits to minimize noise while achieving a desired expression level will require a multi-objective optimization approach.

For high mean expression values we observed a clear benefit of having feedback as compared to constitutive expression. Yet, adding quorum sensing on top of feedback did not decrease noise strength unless the circuit parameters are tuned. That is, the benefit from adding cell-to-cell communication is not structural, but depended on proper choice of the circuit parameters. This result is somewhat counter-intuitive and does not fully agree with previous works reporting a reduction of extrinsic noise in quorum sensing-based gene circuits, e.g. [20], that reported a structural benefit. This may be explained by the different approaches to model extrinsic noise. While we modeled it as parametric variability, most often extrinsic noise has been modeled as an additive stochastic signal essentially analogous to the intrinsic noise term. Thus, if we considered a scenario with intrinsic noise and no extrinsic one while keeping medium-high expression means, our results also showed an important reduction of noise strength when quorum sensing was added to feedback. Though the amount of reduction depended on the circuit parameters, we observed noise reduction for almost any combination of them. Moreover, if we considered additive extrinsic noise, we got qualitatively similar results to the ones when only intrinsic noise was present.

In the hypothetical scenario with no extrinsic noise we also found that adding either feedback or feedback and quorum sensing increased the noise strength with respect to the open loop constitutive gene expression circuit. This result might be explained by the increased complexity introduced by these circuits [91]. Yet, circuit complexity is not the only factor contributing. On the one hand, the circuit with quorum sensing and feedback achieved lower average noise strength values than the less complex only-feedback one in this scenario. On the other, when extrinsic noise was present constitutive expression was clearly noisier than any of the more complex Qs/Fb and NoQs/Fb circuits for high protein mean expression values, though not for low ones where intrinsic noise dominates. Thus, the circuit complexity contribution to noise depends not only on its size, but in the interplay between size and noise structure. Thus, in the medium-high range of mean protein expression, of interest for industrial biotechnology, tuning circuit parameters in the circuit with both quorum sensing and feedback clearly allows coping with both intrinsic noise and extrinsic one, however its structure.

The experimental results, though preliminary, showed a high concordance the computational ones and confirmed the capability of the proposed circuit to reduce noise strength.

## Acknowledgments

Research in this area is partially supported by Spanish government (CICYT DPI2014-55276-C5-1) and European Union (FEDER). Y. Boada thanks grant FPI/2013-3242 of UPV.

## Footnotes

1 Git repository available at https://github.com/incardon

